# Optimization of connectome weights for a neural network model generating both forward and backward locomotion in *C. elegans*

**DOI:** 10.1101/2025.07.21.665845

**Authors:** Taegon Chung, Sangyeol Kim

## Abstract

Previous studies tracking the relationship between manipulations of *C. elegans* neurons and the resulting behavioral changes have called for the development of a connectome-constrained neural network model that describes the cascade from neurons to behavior. However, the model using anatomical connectome weights as they are did not achieve that. Here, we introduce a concept of learning the synaptic weights in our connectome-constrained neural network model based on the leaky-integrator equation while preserving the structural proportions of anatomical synapses. In this process, the weights of gap junctions and chemical synapses in *C. elegans* neurons are optimized. As a result, our neural network model generates plausible *C. elegans* behavior mediated by activity changes in forward and backward command neurons, even without the introduction of pacemaker neurons with intrinsic oscillatory activity. Additionally, we identified necessary or sufficient neurons for maintaining oscillatory patterns on muscular activity that could serve as clues for the central pattern generator in our neural network model. Finally, we provide 10 optimized synaptic weights sets of *C. elegans* that reproduce the results of manipulation experiments on the SMD neurons. This study will facilitate the future study for unraveling the multiscale relationship of “from synapse to behavior” in nervous system.

**Significance Statement:** In the anatomical connectome of *C. elegans*, synaptic weights have been considered less important than network topology in designing neural network models for behavior generation. In this study, we introduce an advanced process that optimizes the synaptic weights of *C. elegans* based on the anatomical connectome weights through machine learning. Our neural network model, incorporating the optimized connectome weights, successfully reproduced both forward and backward locomotion in response to the on/off transitions of command neurons. Through this model, we were able to evaluate how manipulations of specific neurons translate into behavioral changes and to what extent they manifest. This study sets a foundation for exploring the intricate links between neural network structure, neuronal activity, and behavior in living systems.

## Introduction

Reproducing a biological neuronal network of *Caenorhabditis elegans* (*C. elegans*) involves simulating its experimentally observable characteristics. These characteristics fall into two main categories: structural and functional information of its biological neuronal network. The first characteristic, the structure of the neuronal network, also known as the connectome, entails measuring the spatial dimensions of each synapse within the neuronal network and contact volume between neurons. The second characteristic covers the functions of the neuronal network, as reflected in the time-series data of cell activity under various experimental conditions, such as membrane potential or Ca^2+^ activity in neurons and other somatic cells. A model capable of reproducing the structure and functions of the neuronal network could be equivalent to constructing a biological neuronal network. Additionally, it is crucial to elucidate how these structural and functional aspects of neuronal network are interconnected, based on the fundamental principles of neuroscience.

The *C. elegans* is a rod-like animal approximately 1 mm in length, with hermaphroditic adults having 302 neurons, and the transparency of the body facilitates the measurement of cellular Ca^2+^ activity using GCaMP. Neuroscience has long been driven to digitally replicate the neuronal network of *C. elegans*. As a hallmark animal model with fewer neurons than other animal models, such as fruit flies or mice, it presents heightened expectations for successful replication of the neuronal network. Furthermore, *C. elegans* is widely recognized for its importance not only in neuroscience but also throughout the biological sciences. Therefore, there has been an ambition to reverse-engineer the neuronal network of *C. elegans* by creating a digital strategy that operates identically to the real *C. elegans*, aiming to produce a virtual animal for indefinite experimentation without the constraint of mortality.

Previous studies have elucidated the connectome of *C. elegans*, revealing its complete connectome structure, encompassing 302 neurons and 165 somatic cells. This milestone was achieved through the pioneering works of dedicated scientists over recent decades(White et al., 1986; Jarrell et al., 2012; Brittin et al., 2018; Cook et al., 2019). Additionally, there have been further attempts to elucidate the connectome, including studies on variations across life stages(Witvliet et al., 2021) and the Dauer state(Yim et al., 2024), with additional experimental insights(Varshney et al., 2011; Brittin et al., 2021). As revealed by these studies, the connectome has become a crucial determinant for understanding the structure and functions of *C. elegans*’ neuronal circuits, including gap junctions and chemical synapses(White et al., 1986; Hong et al., 2017). Meanwhile, various studies have been conducted to decipher the operation of the neuronal network by measuring the calcium activity in numerous neurons of *C. elegans*(Kato et al., 2015; Nguyen et al., 2016, 2017; Nichols et al., 2017; Kaplan et al., 2020; Hallinen et al., 2021; Susoy et al., 2021; Uzel et al., 2022). Recent research utilizing NeuroPAL technology(Yemini et al., 2021), which facilitates the identification of numerous neurons, has shed light on the functional excitatory-inhibitory relationships among neurons by simultaneously measuring their calcium activities in *C. elegans*(Randi et al., 2023). Several studies have aimed to simulate the neuronal network of *C. elegans* based on the anatomical connectome in which synaptic weights are the structural weights of anatomical contact volume between neurons, proposing various models to explain the neuronal network’s operation for locomotion(Boyle et al., 2012; Kunert et al., 2014, 2017; Izquierdo and Beer, 2018; Sakamoto et al., 2021). However, no study has yet fully succeeded in incorporating the anatomical connectome, neuronal Ca^2+^ activity data, and muscle activity simultaneously into a single model. Previous studies have either solely utilized the network topology of the anatomical connectome(Sakamoto et al., 2021) or employed it with modifications for the repetitive unit structure of a neuronal circuit(Boyle et al., 2012; Izquierdo and Beer, 2018). Some research has criticized the direct usage of anatomical connectome weights on its limitations in explaining the neuronal activities of *C. elegans*(Kaplan et al., 2020; Uzel et al., 2022; Randi et al., 2023). Other research(Kunert et al., 2014, 2017) has incorporated the anatomical connectome(Varshney et al., 2011) as is, generating a central pattern in the simulation. However, they multiplied the projection coefficients of neuronal modes by eigenworm modes(Stephens et al., 2008) to account for muscle cell activities during forward locomotion. This approach, while innovative, was insufficient in fully explaining the translation process from neuronal signals to muscle activities. Most recently, a study aimed at developing a modular neuro-mechanical simulator was conducted in parallel with experiments, comparing the simulated and actual behaviors of *C. elegans* in response to anterior and posterior touch stimuli (Kim et al., 2025). However, the simulation of escape behavior lacked certain dynamic features observed in real organisms—for instance, the simulated touch-stimulus duration was set to over 2 seconds, which is considerably longer than the physiological response time scale of a few dozen milliseconds (Suzuki et al., 2003). In addition, the implementation of proprioception by reversing the direction of the neuromuscular junction pathway, and the application of stimulation not only to sensory neurons but also to interneurons such as AVA and AVB, are physiologically questionable. Furthermore, the simultaneous claims that RIV stimulation induces turning via neuromuscular junctions to ventral neck muscles, and that removal of SMDV completely abolishes the turning behavior, appear logically inconsistent and thus raise confusion in interpretation.

In this study, we present a novel integrative concept and approach that simultaneously accommodates the anatomical connectome, neuronal activity, and muscle activity (Fig. 1) of *C. elegans* and then optimize the synaptic weights of the connectome for gap junction and chemical synapse of *C. elegans* undergoing a forward- and backward-locomotive behavior. By designing and machine-learning of minimizing an error function, we demonstrated that the *C. elegans*’ artificial neuronal network (CANN) model, incorporating the optimized synaptic weights of connectome on the leaky-integrator equation, closely preserves relative synaptic weights of anatomical connectome, as well as SMD neuronal and muscular activity patterns. Furthermore, we confirmed that the body control angle, derived from CANN model and input into the *C. elegans* body mechanical simulator(Chung et al., 2024), accurately reproduced the locomotion pattern of *C. elegans*. Beyond the intended machine-learning objectives of the error function, some machine-learning trajectories of our model similarly revealed the locomotive effects of transient receptor potential cation channel-1,2 (*trp-1,2*) mutants and SMD(D/V) ablation, matching the experimental outcomes(Yeon et al., 2018). Additionally, the neurons that are sufficient or necessary for maintaining the oscillatory activation pattern of body-wall muscle cells were identified in our model. This study serves as a foundation for developing the connectome-constrained neural network model by linking the anatomical connectome, neuronal activities, and muscular activities that induce the behavior of *C. elegans*. It ultimately aims to reverse-engineer the biological neuronal network of *C. elegans* by considering its function, including neuronal activities and various behaviors, and may serve as a new reverse-engineering concept to elucidate the intrinsic working principles of the nervous system.

**Figure 1.**
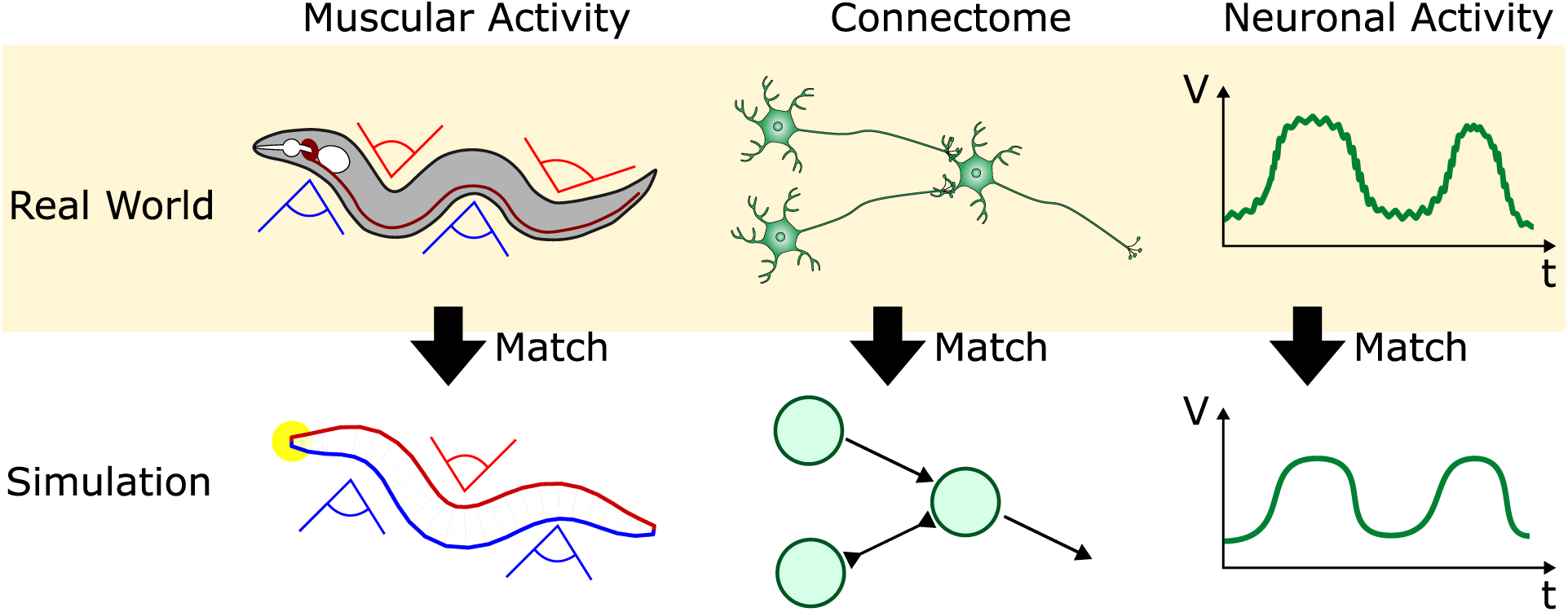
Schematic diagram of the requirements that the neural network model with optimized synaptic weights of *C. elegans* satisfy. The schematic diagram illustrates that the neural network model with optimized synaptic weights is designed to generate muscular activity driving *C. elegans* behavior, adhere as closely as possible to the anatomical neural network topology and synaptic weights, and depict known neural activity. By minimizing the error function to optimize the synaptic weights, we created a CANN (*C. elegans*’ artificial neuronal network) model reproducing muscular activity driving forward and backward locomotion, connectome maintaining the structural proportion of anatomical synapses, and neuronal activity relating the head-bending motion.

## Materials and Methods

### Acquisition and modifications of anatomical connectome data

We obtained the connectome topology and synaptic weights from the anatomical connectome data of Cook et al., 2019. In the anatomical connectome data provided, we utilized information only on the connections between 95 body wall muscles and 162 motor neurons linked to these muscles through synapses, as well as 10 pre-motor interneurons, which are referred to as command neurons for locomotion (Extended Data 1). Thus, we considered a total of 2291 chemical synaptic connections and 626 gap junction connections. The polarity of synapses was set such that the chemical synapses of D-type motor neurons, which correspond to GABAergic neurons, were designated as inhibitory, while those of all other neurons were considered excitatory (Extended Data 2). This study adopts the cell naming convention for *C. elegans* as delineated in previous work(Cook et al., 2019). Given the absence of proprioception information in anatomical connectome data(Cook et al., 2019), we supplemented the CANN model’s motor neurons with connections that facilitate the reception of proprioceptive signals, as detailed in previous studies(Wen et al., 2012; Yeon et al., 2018; Zhan et al., 2023). First, additional chemical synapses were introduced to CANN model between the body wall muscle cells and A/B-type motor neurons. This modification enables the neurons to receive proprioceptive signals from ventral body wall muscle cells. The somas of A/B-type motor neurons, whose locations are detailed in previous work(Haspel and O’Donovan, 2011), are located approximately 0.12 mm apart from the somas of the muscle cells along the anterior-posterior axis. The displacements from the somas of neurons to those of muscle cells are further supported by the findings in previous studies(White et al., 1976; Zhan et al., 2023). Second, new chemical synapses were introduced between the SMD-motor neurons and the four body wall muscle cells (dBWML4, dBWMR4, vBWML4, vBWMR4) in CANN model, enabling SMD motor neurons to receive proprioceptive signals related to the neck-bending angle(Yeon et al., 2018; Kaplan et al., 2020). This modification is supported by previous research, which indicates that the neck-bending angle is in phase with the Ca^2+^ activity in SMDD neurons and antiphase with the Ca^2+^ activity in SMDV neurons(Yeon et al., 2018). To further validate this fact, we employed a feature selection technique(Tibshirani, 1996) on data capturing both dorsoventral body angles and SMD neuron activities simultaneously(Kaplan et al., 2020). This analysis revealed that the variance in SMD neuron Ca^2+^ activity is most significantly associated with the fourth dorsoventral body angle. Consequently, we hypothesized the presence of proprioceptive connections from the fourth body wall muscle cells (dBWML4, dBWMR4, vBWML4, vBWMR4) to SMD neurons, connections that were not previously identified in anatomical studies. Here, we considered a total of 316 proprioceptive connections between neurons and muscles, with the polarity of these connections set as excitatory (Extended Data 2).

### Inference of time constant of *C. elegans* neuron

The time constant is one of the key components that determines the ability of a neuron to compute incoming signals(Lechner et al., 2020). From the experimental data(Liu et al., 2018), we obtained the membrane potential values over time for the RIM, AFD, AIY, and AWA neurons (Fig. S1). To derive a time constant of the membrane potential response to input current for each neuron, we curve-fitted the membrane potential values for the time interval that spans from the initiation of input current application to the point at which the voltage has approached a steady state (1∼1.1 sec) to an exponential function. The time constants were determined to be 0.021 seconds for RIM, 0.017 seconds for AIY, 0.007 seconds for AFD, and 0.018 seconds for AWA. Using interquartile-range-based outlier detection, we classified the time constants of three neurons (RIM, AIY, AWA) as non-outliers. The average time constant of these non-outlier neurons, approximately 0.019 seconds, was then adopted as the representative time constant (𝜏) for all neurons and muscle cells in the CANN model (Fig. S1). The neuron time constants reported vary widely across studies: about 0.1 seconds(Wicks et al., 1996), 0.002 seconds(Boyle et al., 2012), 0.01 seconds(Kunert et al., 2014), and between 0.5 to 2 seconds(Izquierdo and Beer, 2018). Our findings of 0.019 seconds differ from these values. In this paper, we distinguish between ’intra-cellular dynamics,’ the relationship between the current flowing into the cell and its membrane potential excluding synaptic inputs, and ’inter-cellular dynamics,’ the synaptic current from a single synapse determined by the membrane potential of connected cells. Some studies(Wicks et al., 1996; Kunert et al., 2014) have established equations for intra-cellular dynamics based on the capacitance of the phospholipid bilayer and the conductance of leak channels. However, the factors determining the intra-cellular dynamics of a neuron include not only leak channels but also voltage-gated ion channels whose resistance increases or decreases depending on the membrane potential(Hodgkin and Huxley, 1952). Therefore, the use of a leaky-integrator equation for the relation between inflow current and membrane potential means the approximation of complex intra-cellular dynamics into a simpler dynamic where the opening rates of voltage-gated ion channels are fixed and operate like leak channels. Consequently, the methodology employed in prior research(Wicks et al., 1996; Kunert et al., 2014), which derives equations for intra-cellular dynamics exclusively based on membrane capacitance and leak channel conductance, could yield an imprecise time constant in accurately describing actual intra-cellular dynamics. On the other hand, when time constants derived directly from membrane potential measurements over time in current patch clamp experiments(Liu et al., 2018) are applied to a leaky-integrator equation, it allows for a more accurate modeling of the neuron’s intra-cellular dynamics.

### Leaky-integrator equation for membrane potential calculation

The leaky-integrator equation has been used to calculate the membrane potential of each somatic cell in *C. elegans*. This is because the leaky-integrator equation effectively explains the capacitance phenomenon of the cell’s phospholipid bilayer and the ion leakage through leak channels, making it useful for calculating cell membrane potential(Hodgkin and Huxley, 1952). Previous studies have used the leaky-integrator equation to simulate the neuronal network of *C. elegans*. There are slight variations in the leaky-integrator equation among previous studies(Hodgkin and Huxley, 1952; Lockery and Sejnowski, 1992; Wicks et al., 1996; Boyle et al., 2012; Kunert et al., 2014; Izquierdo and Beer, 2018; Sakamoto et al., 2021). We have also developed a version of the leaky-integrator similar to those found in prior research (Details in “Formulating the leaky-integrator equation” of Supplementary Materials). The leaky-integrator equation used in this paper is as follows:

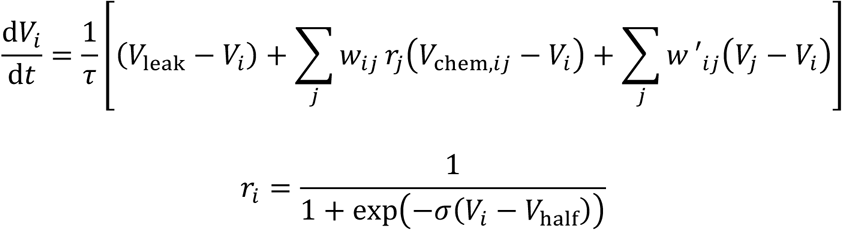

where 𝜎 = 5.9, 𝑉_half_ = 0.5. 𝑡 represents time. 𝑉 represents the normalized membrane potential, a value between 0 and 1 in this study (see “Definition of normalized membrane potential” of Supplementary Materials). Specifically, 𝑉_*i*_ represents the membrane potential of cell 𝑖, 𝑉_leak_ (=0) represents the electrical potential difference due to the membrane’s leak channel, and 𝑉_chem,*ij*_ represents the electrical potential difference due to the ion channel of the chemical synapse from presynaptic neuron cell 𝑗 to postsynaptic cell 𝑖. For an excitatory chemical synapse, 𝑉_chem,*ij*_ equals 1; for an inhibitory one, it equals 0. 𝜏 represents the time constant. The parameter 𝑤_*ij*_ denotes the synaptic weight of the chemical synapse from cell 𝑗 to cell 𝑖 (𝑤_*ij*_ ≥ 0), whereas 𝑤′_*ij*_ denotes the synaptic weight of the gap junction between cell 𝑖 and cell 𝑗 (𝑤′_*ij*_ ≥ 0). The parameters 𝑤_*ij*_and 𝑤′_*ij*_ are constants that are independent of time 𝑡. The variable 𝑟_*i*_represents the ion channel opening rate of the chemical synapse from cell 𝑖; when cell 𝑖 is a muscle cell, 𝑟_*i*_also indicates the degree of muscle contraction. In reality, the degree of muscle contraction depends on the frequency of muscle action potentials (Gao and Zhen, 2011; Liu et al., 2011b, 2011a), but this simplified abstraction is adopted in our model. There are lower and upper limits for both ion channel opening rate and degree of muscle contraction (𝑟_*i*_ ∈ (0,1)).

### Configuration of the membrane potential of command neurons

As in previous research(Sakamoto et al., 2021), we ensured that the forward-locomotion-command-neurons (AVB, PVC) and the backward-locomotion-command-neurons (AVA, AVD, AVE) alternately had opposing membrane potentials(Chalfie et al., 1985; Riddle et al., 1997; Kato et al., 2015; Uzel et al., 2022; Roberts et al., 2016). A forward-locomotion-command is a state in which the membrane potentials of forward-locomotion-command-neurons are 1, and those of backward-locomotion-command-neurons are 0. Conversely, a backward-locomotion-command is a state in which the membrane potentials of forward-locomotion-command-neurons are 0, and those of backward-locomotion-command-neurons are 1. When solving the leaky-integrator equation, the command at each time point 𝑡 was determined to be either a forward-locomotion-command or a backward-locomotion-command.

### Numerical method for solution of leaky-integrator equation

As the complexity of the neuronal network increases, the leaky-integrator equation becomes increasingly susceptible to stiffness issues of numerical integration(Lechner et al., 2018). We employed the semi-implicit Euler method for numerical integration to resolve the stability issues of solving ordinary differential equations(Lechner et al., 2018, 2020). To correct inaccuracies in the decay rate of membrane potential, we employed a modified version of the semi-implicit Euler method with a time step (𝛥𝑡) of 0.01 sec (Details in “Modified semi-implicit Euler method” in Supplementary Materials).

### Definition of the error function

Inspired by research on methods for optimizing multiple objective functions simultaneously(Peng et al., 2018), we designed the combined form of the error function 𝐸 to ensure that the CANN model could explain a variety of *C. elegans* experimental data(Vidal-Gadea et al., 2011; Yeon et al., 2018; Cook et al., 2019). By minimizing the error function 𝐸, we obtained the optimized synaptic weights 𝑤_*ij*_ and 𝑤′_*ij*_, as well as the initial membrane potential values 𝑉^(0)^_*i*_ (∗^(*t*)^: value of a time-dependent variable at time 𝑡) for each neuron.

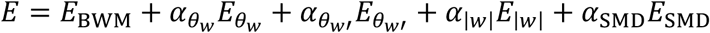

𝛼 denotes the coefficient of each error term (Details in “Details of the definition of the error function “ of Supplementary Materials).

### Update process of synaptic weights for minimization of the error function

After appropriately adjusting the 𝛼 values (𝛼_θ*w*_ = 0.5, 𝛼_θ*w*_ = 4, 𝛼_|5|_ = 0.1, 𝛼_627_ = 1), the minimization of the error function 𝐸 effectively reduces all its error terms. This process enables the identification of parameters 𝐰 and 𝐰′, allowing the CANN model to explain a wide range of distinct wet experimental data (Fig. 1). The steps to find the 𝑤_*ij*_ and 𝑤′_*ij*_ values that minimize the error function 𝐸 are as follows. First, 𝑤_*ij*_ and 𝑤′_*ij*_ were initialized with random values, where 𝑤_*ij*_ ranges from 0.001 to 2, and 𝑤′_*ij*_from 0.001 to 1. 𝑉_*i*_ at 𝑡 = 0 was initialized as 𝑉^(-)^. The leaky-integrator equation was numerically solved to calculate 𝑉_*i*_and 𝑟_*i*_ over time 𝑡 (see “Numerical integration of leaky-integrator equation” of Supplementary Materials). The value of the error function 𝐸 was computed using the values of 𝑉_*i*_ and 𝑟_*i*_ over time 𝑡, along with 𝑤_*ij*_ and 𝑤′_*ij*_. Adam optimizer(Kingma and Ba, 2014) was employed to update the values of 𝑤_*ij*_, 𝑤′_*ij*_, and 𝑉^(-)^ based on the derivatives of 𝐸 with respect to 𝑤_*ij*_, 𝑤′_*ij*_, or 𝑉^(-)^ (see “Derivatives of the error function” of Supplementary Materials) to minimize the error function 𝐸, with the Adam step size set to 0.001, and the Adam parameters 𝛽_1_ to 0.9 and 𝛽_2_ to 0.999. We created and utilized an adaptive error minimization method to reduce the computational cost of the error minimization process (see “Adaptive error minimization” of Supplementary Materials).

### Stepwise neuron elimination

Stepwise neuron elimination is a method for identifying the sufficient neuron set required to maintain the oscillatory activation pattern of body-wall muscle cells for forward or backward locomotion. During locomotion generation in the CANN model, neurons are sequentially ablated one at a time, starting with those that have the least impact on the oscillatory activation pattern of the body-wall muscle cells. Intentional simultaneous ablation of left–right neuron pairs was not performed. The method is as follows (Fig. S2A). First, a base network model was set as a CANN model where all neurons were intact. Then, the following steps were repeated. Copies of the base network model were created, with each copy having one neuron removed. The membrane potential (𝑉_*i*ϵBWM_^(*t*^)) of the body muscle membrane was computed for each modified network model under the forward-locomotion-command (Fig. S2B). The dynamic mode decomposition (DMD)(Kutz et al., 2016) was applied to the membrane potential data for each of the 95 muscles within the 1 to 10-second time range. For efficient computation, prior to DMD application, we subtracted the time-averaged membrane potential from the data of each muscle. By examining the absolute value of the eigenvalue of the first DMD mode and the initial coefficient at 𝑡 = 1 sec, we identified the ablation scenario where the coefficient magnitude of the first DMD mode at 𝑡 = 4 sec (≡ 𝛾) was closest to 1. This established the base network model for subsequent steps. The persistence of muscle membrane potential oscillation is directly related to the absolute value of the eigenvalue from the first DMD mode; the closer this value is to 1, the longer the oscillation persists. The process was repeated until all motor neurons are removed in the CANN model. Next, if 𝛾 exceeds 0.1 and the period of the first DMD mode (≡ 𝑇) ranges from 0 to 4.5 seconds, with the fewest unablated motor neurons, we defined that step’s motor neuron group as the sufficient neuron group.

### Single neuron elimination

To identify motor neuron candidates that are necessary for oscillatory patterns maintained in the membrane potential of body muscle cells for forward and backward locomotion, we analyzed necessary neurons for each machine-learning trajectory. A necessary neuron is a motor neuron within the CANN model that, when ablated in isolation, leads to the oscillation disappearance of membrane potential of the 95 muscles. The criterion for oscillation disappearance is determined by |𝛾| and 𝑇. Oscillation is considered to have disappeared when |𝛾| < 0.1, 𝑇 ≤ 0 sec, or 𝑇 ≥ 4.5 sec.

### Analysis Code and Tools

The analyses were conducted using Python(van Rossum, 1995), NumPy(Harris et al., 2020), and Numba(Lam et al., 2015). Visualizations were created with Matplotlib(Hunter, 2007). The source code for these analyses will be made available upon publication of the paper or sooner if requested by a reviewer or editor.

## Results

### Synaptic weights of gap junctions and chemical synapses were optimized with high correlations to anatomical connectomes

We developed a CANN model that controls *C. elegans* muscle cells for locomotion using a leaky-integrator equation (Methods). Muscle contraction is necessary for the worm to locomote. The degree of muscle contraction (𝑟_*i*_, 𝑖 ∈ {BWM}) is determined by the membrane potential of a muscle cell (𝑉_*i*_, 𝑖 ∈ {BWM}). The time derivative of the membrane potential of a muscle cell is dependent on synaptic currents. For the CANN model to generate appropriate synaptic currents, 3233 synaptic weight values (𝑤_*ij*_and 𝑤′_*ij*_) of connectomes must be accurately adjusted. We have developed a neuronal network model that generates the muscle membrane potentials necessary for locomotion by formulating an error function and minimizing the error function by adjusting synaptic weights (Methods). The membrane potentials of each cell at time 𝑡 = 0 (𝑉^(-)^), necessary for *C. elegans* to initiate locomotion, were also determined using the same method. This approach was based on the assumption that the probability of initiating locomotion would be low if *C. elegans* starts from an improper state, such as a sleeping state(Nichols et al., 2017).

We ran 300 machine-learning trajectories, of which 57 resulted in a CANN model that produced the muscle membrane potentials necessary for forward and backward locomotion. In other words, all 57 machine-learning trajectories successfully converged the value of our designed error function below a certain threshold (Extended Data 3). Therefore, we performed further analysis only on those 57 machine-learning trajectories. For each of these machine-learning trajectories, we calculated a correlation between optimized synaptic weights and anatomical synaptic weights of chemical synapses (Fig. 2A), and the median correlation among the machine-learning trajectories was 0.91 (Interquartile range, IQR: 0.01). Similarly, for each machine-learning trajectory, a correlation between optimized synaptic weights and anatomical synaptic weights of gap junctions were calculated (Fig. 2B), and the median correlation among the machine-learning trajectories was 0.85 (IQR: 0.04). This implies that most of the relative ratios among anatomical synaptic weights are preserved within optimized synaptic weights derived from these machine-learning trajectories.

**Figure 2.**
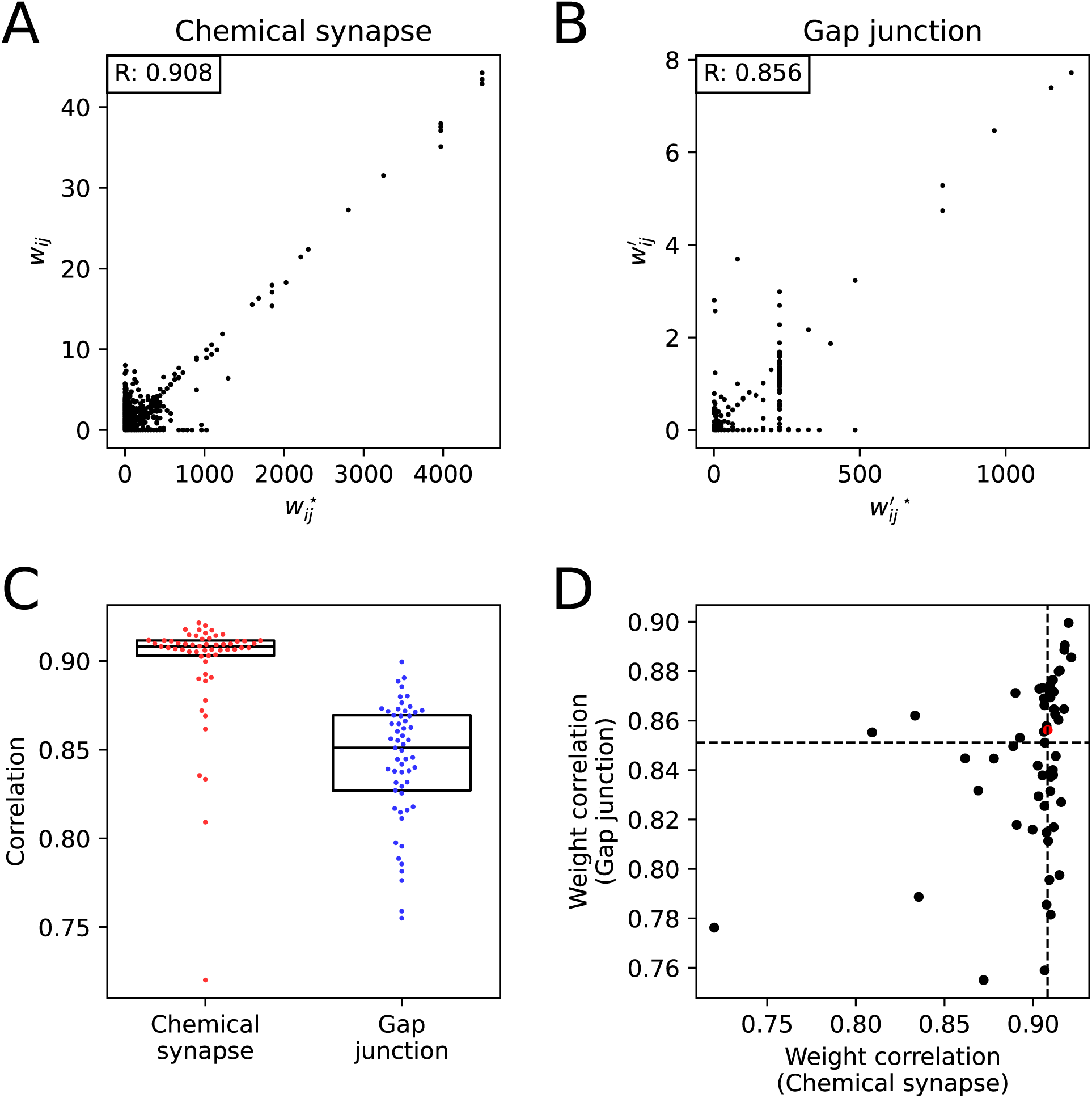
Synaptic weights comparison between anatomical connectome and CANN model. **(A, B)** Scatter plots comparing anatomical synaptic weights to optimized synaptic weights of a representative machine-learning trajectory. The number in the top left corner of each box indicates the correlation coefficient (R). **(A)** Chemical Synapse **(B)** Gap junction. **(C)** Correlation value between anatomical synaptic weights and optimized synaptic weights for each machine-learning trajectory. Each point represents a machine-learning trajectory (57 in total for each, red and blue). The y-axis component of each point indicates the correlation value (red for chemical synapse, blue for gap junction). The x-axis component of each point does not represent any significant value; it is determined solely to minimize overlap between points. The central horizontal line in the black box represents the median (Q2), the top line represents the first quartile (Q1), and the bottom line represents the third quartile (Q3). **(D)** Scatter plot of chemical synapse weight correlation and gap junction weight correlation for each machine-learning trajectory. Dotted lines represent the median value for each axis, and the red point denotes the representative machine-learning trajectory of (A, B).

The correlation values between optimized synaptic weights and anatomical synaptic weights of chemical synapses showed a concentrated distribution with an IQR of 0.01, and those for gap junctions displayed a more dispersed distribution than that of the chemical synapse with an IQR of 0.04 (Fig. 2C). There was no clear relationship between gap junction weight correlations and chemical synapse weight correlations in CANN model (Fig. 2D).

### Spatio-temporal track of body, and SMD neurons activities, of *C. elegans* during locomotion were reproduced using CANN model with optimized synaptic weights

Degrees of muscle contraction (𝑟_*i*_, 𝑖 ∈ {BWM}) were calculated from a CANN model of each machine-learning trajectory under forward-locomotion-command for 10 seconds of simulation time. The body control angles (𝜃_’;<”,=_, 𝑘 ∈ {1, 2, …, 24}) were derived from these degrees of muscle contraction (Details in “Method to calculate body control angle” of Supplementary Materials) and exhibited a sine wave pattern, resembling the kymogram pattern observed in real *C. elegans* (Fig. 3A). Inputting the body control angles into the mechanical body simulator (hereafter referred to as ElegansBot)(Chung et al., 2024), ElegansBot exhibited spatio-temporal track of forward locomotion (Fig. 3B, Extended Data 4). ElegansBot is a two-dimensional rigid body chain model that simulates *C. elegans* locomotion by applying Newtonian equations of motion to each body segment. It reproduces various locomotive behaviors and quantifies force distributions, enabling the study of muscle-neuron interactions and environmental friction effects. Similarly, degrees of muscle contraction were calculated from a CANN model under backward-locomotion-command, and the body control angles were calculated from these degrees of muscle contraction and exhibited a sine wave pattern of backward locomotion kymogram (Fig. 3C). Inputting the body control angles into the ElegansBot, ElegansBot showed backward locomotion track (Fig. 3D, Extended Data 4). From body control angles of forward and backward locomotion, period and linear wavenumber of locomotion were derived for each machine-learning trajectory, by DMD analysis on the body control angles and sine-function curve-fitting to the oscillating DMD mode pair of a longest period. (Fig. 3E, 3F). From the spatio-temporal track of locomotion, the mechanical speed was calculated for each machine-learning trajectory (Fig. 3G). The median period, the median linear wavenumber, and the median speed across machine-learning trajectories were 1.61 sec (IQR: 0.04 sec), 1.80 (IQR: 0.03), and 0.21 mm/sec (IQR: 0.01 mm/sec) for forward locomotion and 1.63 sec (IQR: 0.06 sec), 1.83 (IQR: 0.05), and 0.20 mm/sec (IQR: 0.02 mm/sec) for backward locomotion. When a real *C. elegans* crawls to forward on an agar plate, the period, wavenumber, and speed are 1.6 sec, 1.83, and 0.21 mm/sec, respectively(Chung et al., 2024). This indicates that the CANN model of each machine-learning trajectory can generate control signals necessary for both forward and backward locomotion. Under a command alternating between forward-locomotion-command and backward-locomotion-command every 3.2 seconds, with a total duration of 10 seconds, degrees of muscle contraction were generated from CANN model. The body control angles, calculated from the degrees of muscle contraction, showed temporally alternating forward and backward locomotion patterns. To check whether membrane potential of SMD neurons calculated from CANN model and neck-bending angle has agonistic or antagonistic relationship, we compared the membrane potential of SMD neurons and the neck-bending angle (𝜃_’;<”,>_) for each machine-learning trajectory (Fig. 3H, Extended Data 4) by calculating Pearson correlation between them (Fig. 3I). The median correlation among machine-learning trajectories was 0.98 (IQR: 0.02) for SMDDL, 0.96 (IQR: 0.07) for SMDDR, -0.8 (IQR: 0.2) for SMDVL, -0.9 (IQR: 0.2) for SMDVR. These results were consistent with the wet experiments that showed that the Ca^2+^ activity of the SMDD neuron was in phase with the neck-bending angle, and the Ca^2+^ activity of the SMDV neuron was in antiphase with the neck-bending angle(Yeon et al., 2018; Kaplan et al., 2020).

**Figure 3.**
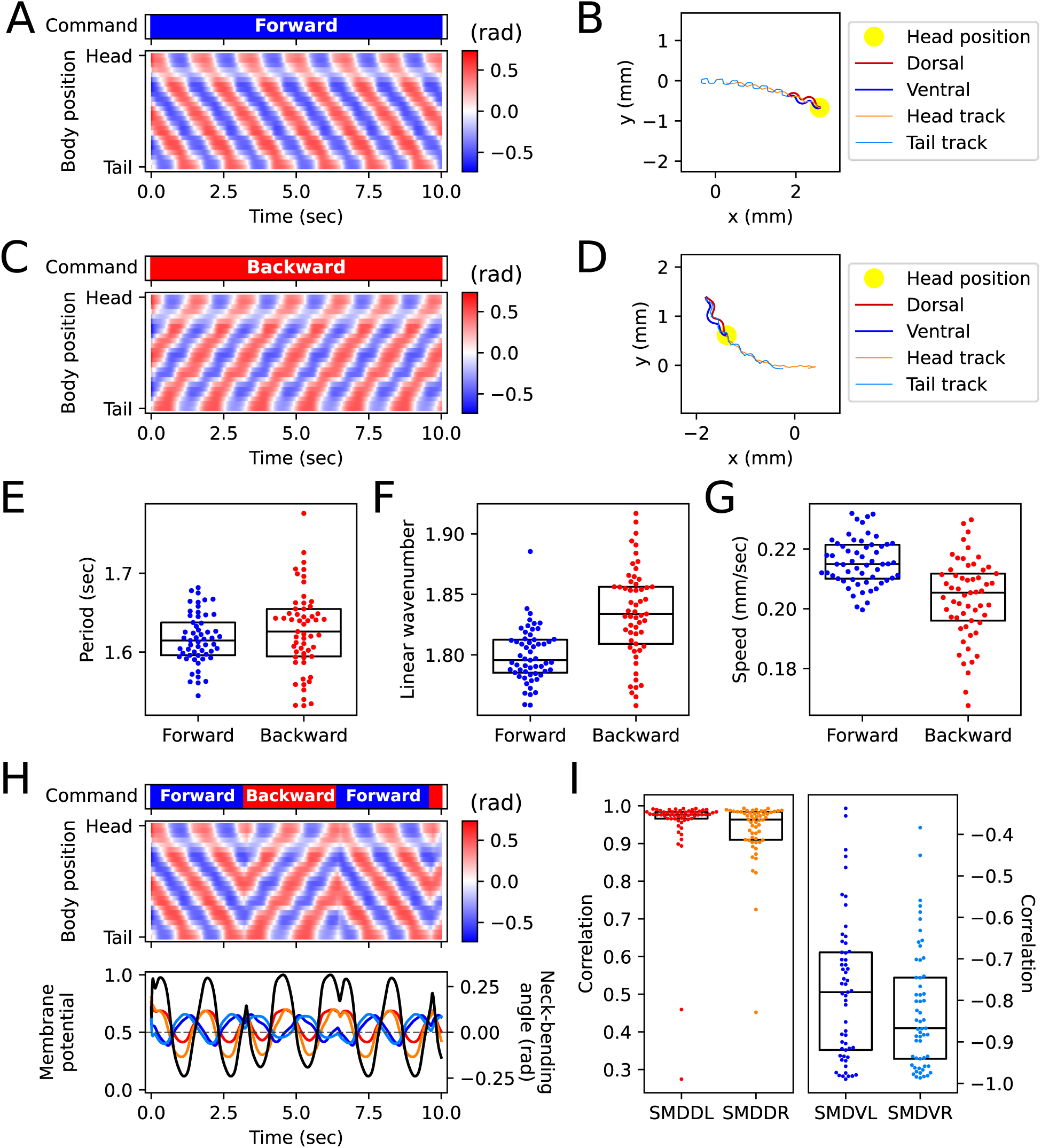
Neuronal and muscular activity reproduced using CANN model. **(A)** Body control angles determined by the muscle membrane potential over time from CANN model of the representative machine-learning trajectory under forward-locomotion-command. Upper: Locomotion command (blue: forward-locomotion-command, red: backward-locomotion-command). Lower: Body control angles. **(B)** The ElegansBot locomotion track derived from the input body control angles of (A). The yellow circle indicates the head position of *C. elegans*. The orange and sky-blue lines mean head and tail tracks, respectively. **(C)** Body control angles over time determined by the muscle membrane potentials from CANN model of the representative machine-learning trajectory under backward-locomotion-command. **(D)** The ElegansBot locomotion track derived from the input body control angles of (C). **(E)** Oscillation period calculated from body control angles. Each point represents a machine-learning trajectory (57 in total for each, blue and red). The y-axis component of each point indicates the correlation value (blue for forward locomotion, red for backward locomotion). The x-axis component of each point does not convey any meaning; it is determined solely to minimize overlap between points. The central horizontal line in the black box represents the median (Q2), the top line represents the first quartile (Q1), and the bottom line represents the third quartile (Q3). **(F)** Oscillation linear wavenumber calculated from body control angles. **(G)** ElegansBot locomotion speed resulted from input body control angles of each machine-learning trajectory. **(H)** Locomotion command (upper panel) triggered the CANN model to generate body control angles (middle panel), the membrane potential of SMD neurons (lower panel: red for SMDDL, orange for SMDDR, blue for SMDVL, sky-blue for SMDVR), and neck-bending angle (black), resulted from the representative machine-learning trajectory. **(I)** Pearson correlation between neck-bending angle and membrane potential of SMD neurons. Each point represents a machine-learning trajectory (57 in total for each, red, orange, blue, and sky-blue).

In summary, we confirmed that the CANN model, the optimized connectomes derived in this study, exhibited forward and backward locomotion functions and SMD neurons’ membrane patterns similar to those observed in the experiment. Thus, we have developed a CANN model that meets all our targeted objectives (Fig. 1). We further confirmed the CANN model’s capability to generate locomotion by integrating it with ElegansBot and observing the spatio-temporal track of mechanical locomotion.

### The sufficient and necessary neurons that sustain the periodic activation of *C. elegans* body-wall muscle cells have been identified

*C. elegans* produces sinusoidal wave patterns during locomotion. For such sinusoidal wave patterns to exist, a central pattern generator (CPG) that produces oscillatory signals must be present within *C. elegans*. Previous studies(Wen et al., 2012; Fouad et al., 2018; Gao et al., 2018; Xu et al., 2018) have provided hypotheses and experimental evidence that the CPG of *C. elegans* resides within the neuronal circuit and is facilitated by proprioception. In this study, we present results identifying the neuron groups that are sufficient or necessary for sustaining the oscillatory activation of muscle cells, which may provide clues about the *C. elegans* CPG. This was achieved using ablation-based methods, including stepwise neuron elimination or single neuron elimination (Methods).

The sufficient neuron group identified through stepwise neuron elimination represents the minimal set of motor neurons predicted to be necessary for generating muscle membrane potential oscillation. The necessary neuron is a neuron whose removal from an intact CANN model eliminates muscle membrane potential oscillation. Among the 57 machine-learning trajectories that have learned both forward and backward locomotion, differences were observed in the neuron configuration of the sufficient neuron group. Consequently, we counted the number of times each neuron appeared as a member of the sufficient neuron group across the 57 machine-learning trajectories. Similarly, we counted how often each neuron appeared as a necessary neuron across the 57 machine-learning trajectories. Neurons that appear more frequently can be considered more crucial for generating oscillatory membrane potentials in muscles within the CANN model.

The forward locomotion sufficient neuron and forward locomotion necessary neuron identified through these two methods are similar in that their counts are comparable (Fig. 4A). Excluding neurons with both sufficient neuron count and necessary neuron count for forward locomotion under 10 out of 57, the Pearson correlation between sufficient neuron counts and necessary neuron counts for forward locomotion was 0.88 (0.94 when including the excluded neurons). A strong correlation between the two counts indicates that neurons with high counts are both necessary and sufficient for maintaining the oscillatory membrane potential pattern in muscles within the CANN model. Similarly to the forward locomotion scenario, when excluding neurons with both sufficient neuron count and necessary neuron count for backward locomotion under 10 out of 57, the Pearson correlation between sufficient neuron counts and necessary neuron counts for backward locomotion was found to be 0.83 (0.96 when including the excluded neurons) (Fig. 4B).

**Figure 4.**
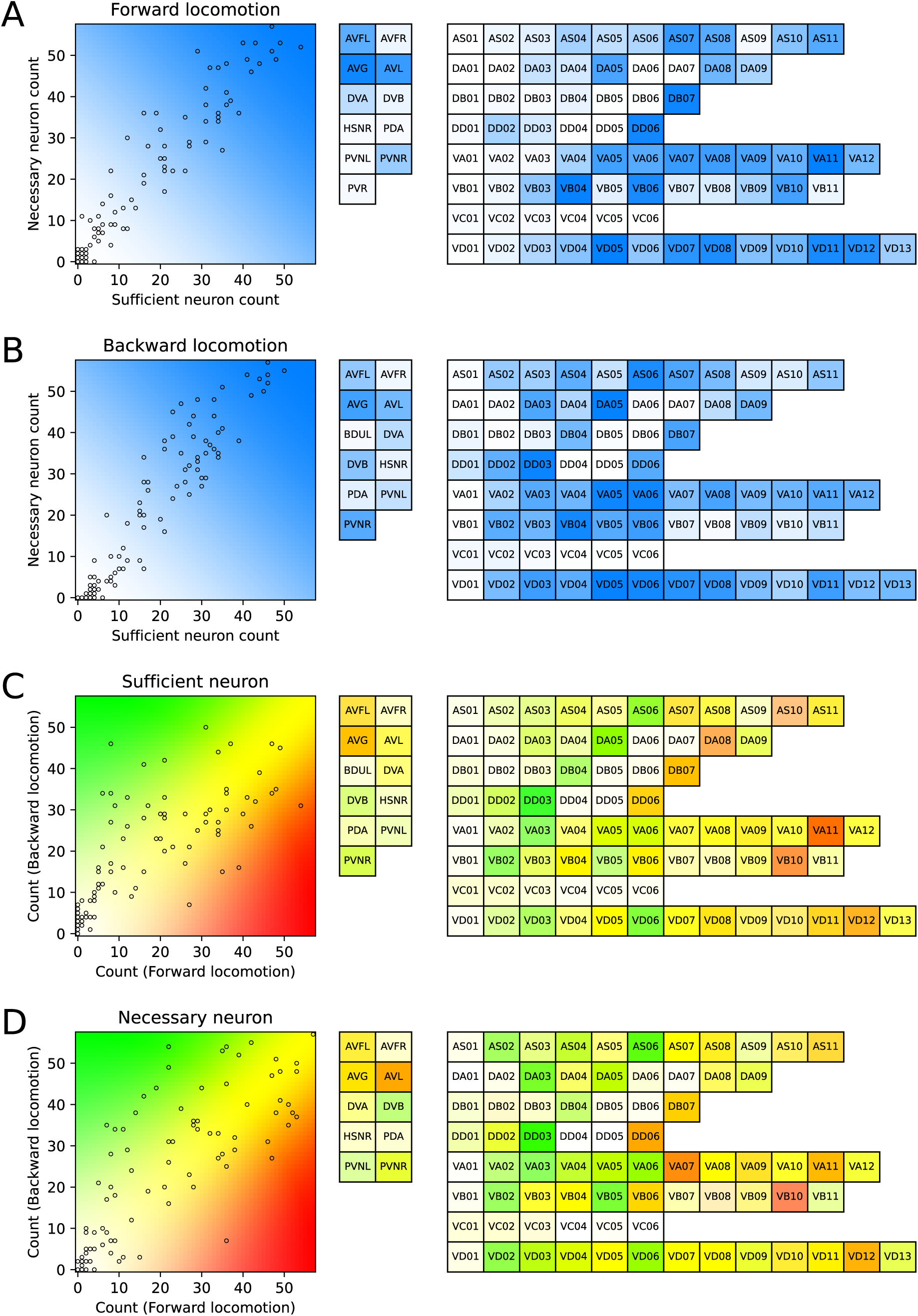
Identification of sufficient neuron sets or necessary neurons for maintaining oscillatory patterns on muscular activity. **(A)** Sufficient neuron counts versus necessary neuron counts for forward locomotion. Each hollow dot in the left diagram represents a motor neuron. The x-coordinate of the hollow dot indicates the number of machine-learning trajectories for which the motor neuron is selected as a sufficient neuron, and the y-coordinate indicates the number of trajectories for which the motor neuron is selected as a necessary neuron for forward locomotion. The text in each cell of the right table indicates the neuron’s name, and the color corresponds to the background color at the location of the corresponding hollow dot in the left diagram. If different motor neurons have the same sufficient neuron counts and the same necessary neuron counts, the corresponding hollow dots for the motor neurons in the left diagram overlap. **(B)** Sufficient neuron counts versus necessary neuron counts for backward locomotion. This follows the same representation and methodology as described for forward locomotion in (A). **(C)** Forward locomotion versus backward locomotion sufficient neuron counts. This follows the same representation and methodology as described in (A). A motor neuron in the table is more important to the forward locomotion if the color is redder, more important to the backward locomotion if the color is greener, and equally important to both forward and backward locomotion if the color is yellow. **(D)** Forward locomotion versus backward locomotion necessary neuron counts. This follows the same representation and methodology as described in (C).

We also compared the sufficient neuron counts between forward and backward locomotion (Fig. 4C). When excluding neurons with sufficient neuron counts under 10 out of 57 for both forward and backward locomotion, the correlation between sufficient neuron counts for forward and backward locomotion was 0.46 (0.82 when including the excluded neurons). A relatively weak correlation between the two counts suggests that the composition of neurons maintaining the oscillatory membrane potential pattern in muscles within the CANN model may differ between forward and backward locomotion. The average number of sufficient neurons was 27.7 for forward locomotion and 34.8 for backward locomotion, respectively. A similar relationship was observed in the necessary neuron counts. When excluding neurons with necessary neuron counts under 10 out of 57 for both forward and backward locomotion, the correlation between necessary neuron counts for forward and backward locomotion was 0.52 (0.78 when including the excluded neurons) (Fig. 4D). The average number of necessary neurons was 33.7 for forward locomotion and 38.7 for backward locomotion, respectively.

### The impact of *trp-1,2* mutation and SMD(D/V) ablation observed in the proprioception experiments of *C. elegans* was explained and reproduced

To assess whether our CANN model reproduces the results of proprioception experiments from prior research(Yeon et al., 2018)(Fig. 5A, 5B), we calculated the body control angle from the muscle cell contraction generated by the CANN model under forward-locomotion-command for a duration of 10 seconds and computed the locomotion track of *C. elegans* by inputting the body control angle into the ElegansBot(Chung et al., 2024) (Fig. 5A). For each of the 57 machine-learning trajectories which can perform both forward and backward locomotion, we calculated the degrees of muscle contraction from CANN model under forward-locomotion-command and then calculated the body control angles based on these degrees of muscle contraction. We calculated the average body control angle over all time and angle index pair (𝑡, 𝑖). If the absolute value of this average was above 0.01, it might not reveal the changes in locomotion caused by mutation or ablation, as each wild-type *C. elegans* of these machine-learning trajectories performs circular locomotion. Therefore, we excluded 11 machine-learning trajectories with the absolute value of an average body control angle above 0.01 and considered the remaining 46 machine-learning trajectories as participant trajectories for reproducing the results of the wet experiments.

**Figure 5.**
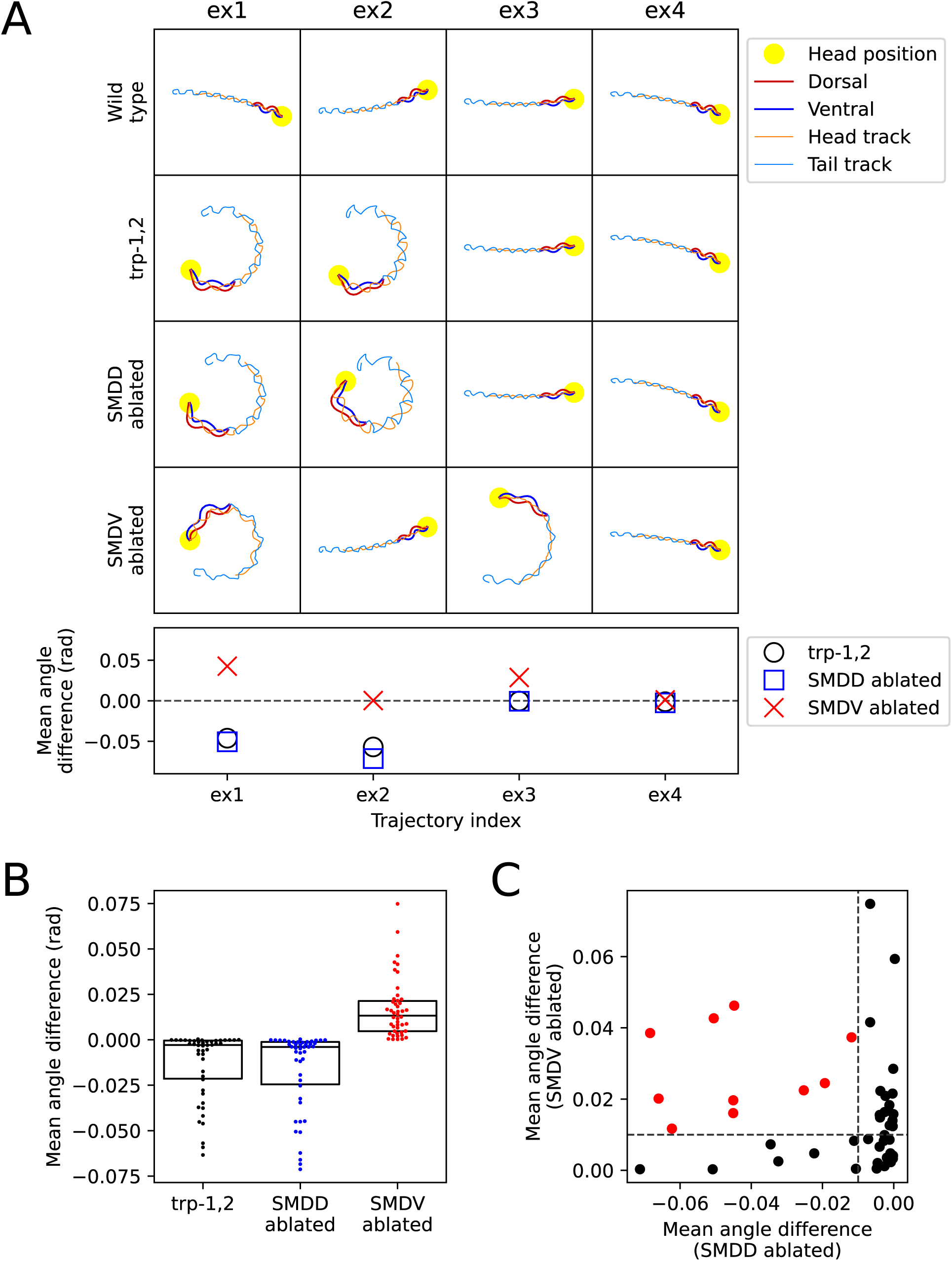
Implementation of proprioceptive sensory restriction and neuronal ablation experiments using CANN model. **(A)** The impact of *trp-1,2* mutation (elimination of proprioceptive sensations of SMDD neuron) and SMD(D/V) neuron ablation on the locomotion track of the worm for each of four example machine-learning trajectories (ex1, ex2, ex3, ex4, each representing the index of a trajectory). Top: The locomotion track of each example machine-learning trajectory. Bottom: Mean angle difference. **(B)** Distribution of mean angle differences of participant trajectories. Each point represents a participant trajectory (46 in total for each, black, blue, and red). The central line in the box plot represents the median (Q2), the top edge of the box represents the first quartile (Q1), and the bottom edge represents the third quartile (Q3). **(C)** Scatter plot of the mean angle difference for each participant trajectory. Red dots represent participant trajectories that match the locomotion phenotype caused by *trp-1,2* mutation and SMD(D/V) ablation, as reported in the wet experiment paper(Yeon et al., 2018).

For each of the 46 participant trajectories, we compared how locomotion differed between the unmanipulated case (wild-type) and the manipulated case (*trp-1,2*, SMD(D/V) ablation). We calculated the difference in body control angle by subtracting the body control angles of the unmanipulated case from that of the manipulated case. We calculated the average of the difference in body control angle over time and angle index pair (𝑡, 𝑖), naming this as the mean angle difference (Fig. 5B). We defined the manipulated case as performing ventrally biased locomotion if the mean angle difference was below -0.01, and as performing dorsally biased locomotion if the mean angle difference was above 0.01.

The TRP-1,2 knockout mutant (*trp-1,2*) is known to exhibit ventrally biased locomotion. This is because the channel proteins expressed by the TRP-1,2 genes are involved in the neck-bending angle proprioception of SMDD neurons, and this proprioception is impaired by the *trp-1,2* mutation(Yeon et al., 2018). To mimic the effects of this mutation, the proprioceptive connection between SMDD neurons and muscles was removed in the CANN model, blocking the proprioception signal. As a result, 15 out of the 46 participant trajectories exhibited the ventrally biased locomotion (Fig. 5B).

When SMDD (or SMDV) neurons in actual *C. elegans* are ablated, it was known to result in ventrally-biased (or dorsally-biased) locomotion(Yeon et al., 2018). To mimic the ablation of specific neurons in actual *C. elegans* within the CANN model, all synapses connected to the respective neurons were removed. We termed this action of removing all synapses connected to specific neurons in the CANN model as ’neuronal ablation’ in the simulation. In wet experiments, ’ablation’ refers to the destruction of specific neurons using lasers or cautery. For the total of 46 participant trajectories, 17 trajectories exhibited dorsally biased locomotion when SMDD was ablated, and 26 trajectories exhibited ventrally biased locomotion when SMDV was ablated (Fig. 5B). Notably, 10 trajectories demonstrated both effects, showing the dorsally biased locomotion when SMDD was ablated and the ventrally biased locomotion when SMDV was ablated (Fig. 5C). These 10 trajectories also reproduced the effect of the *trp-1,2* mutation (Extended Data 5). In other words, since these 10 trajectories out of a total of 57 successfully reproduced the effects of the *trp-1,2* mutation and SMD(D/V) ablation, we evaluated the optimized synaptic weight sets of these 10 trajectories as the most promising synaptic weight data (Extended Data 6). When each of these 10 most promising synaptic weight sets is applied to the CANN model, the membrane potentials of all cells included in the model can be observed during alternating forward and backward locomotion (Extended Data 7).

## Discussion

### CANN model, the optimized weights of connectome, explained and reproduced various experimental results of *C. elegans*

Previously, no consistent method existed for simultaneously reproducing various types of experimental data of *C. elegans*. However, some studies successfully reproduced individual types of experiments, each with its unique advantages and disadvantages(Kunert et al., 2017; Izquierdo and Beer, 2018; Sakamoto et al., 2021). One model managed to perfectly replicate the topology and partially reproduce the kymogram, but the reproduction of anatomical synaptic weights of the connectome was incomplete(Sakamoto et al., 2021). Another model successfully reproduced the crawling kymogram, though it determined the connectome’s topology arbitrarily, and the synaptic weights of the connectome were identified through computational search methods, leaving no guarantee or indication that the modeled synaptic weights would match anatomical synaptic weights (Izquierdo and Beer, 2018). Moreover, the model had both excitatory and inhibitory synapses from a single SMD (or RMD) motor neuron. Yet another model used the anatomical connectome as is(Kunert et al., 2017). However, the equation determining muscle activity to exhibit a sinusoidal pattern, by multiplying projection coefficients of neuronal modes to eigenmodes(Stephens et al., 2008), was qualitatively different from the equation governing neuronal activity in the model. Thus, strategies focused on perfectly reproducing a single type of wet experiments may prove to be impractical. Instead, we adapted the error function of the previous model(Sakamoto et al., 2021) to train it to simulate not only muscle activity patterns but also the connectome’s synaptic weights and neuronal activity patterns. Consequently, our strategy does not solely focus on perfectly reproducing one of anatomical connectome topology, connectome’s synaptic weight, muscular activity, or neuronal activity, but aims to adequately reproduce all four of them. However, setting a high value of coefficient on a single error term in our model’s error function can lead to explaining one type of experiment well while failing to account for others, similar to previous studies(Kunert et al., 2017; Izquierdo and Beer, 2018; Sakamoto et al., 2021). This is due to the difficulty in reducing all error terms to the desired level, as increasing the coefficient of one error term to reduce its value can result in a trade-off where other error terms increase. Nonetheless, we have demonstrated the effectiveness and practicality of our methodology by appropriately reproducing anatomical connectome, muscular activity, and neuronal activity within a single model.

The CANN model, optimized using this strategy, demonstrates the ability to generate both forward and backward locomotion in *C. elegans* without the need for a preferred pacemaker neuron. In neural network models incorporating pacemaker neurons, these neurons autonomously generate oscillatory activation patterns, and the network outputs appropriate muscle contraction/relaxation patterns to produce forward and backward locomotion based on these intrinsic oscillations. However, there is still little evidence supporting the existence of pacemaker neurons specifically responsible for both forward and backward locomotion in *C. elegans*. Instead, it is well established that the on/off activity of certain command neurons, such as AVA and AVB, triggers forward and backward locomotion. The CANN model captures the function of these command neurons and demonstrates that reversing their activity can switch between forward and backward movement. The oscillatory membrane potential patterns generated in the CANN model rely heavily on proprioceptive connections that are organized and operated across A/B-type motor neurons and SMD neurons. The feedback loop between these neurons and muscle cells enables the generation of such oscillatory patterns. Furthermore, in the 10 most promising CANN models that successfully replicated the effects of SMD(D/V) neuron ablation experiments, we observed that the removal of proprioceptive connection in SMD neurons (simulating the *trp-1,2* mutation experiment) alone was sufficient to disrupt normal forward locomotion, leading to circular locomotion. This suggests that proprioceptive connections are essential for maintaining the normal oscillatory pattern of muscle membrane potentials.

### Application of our strategy to other nervous system

Our new strategy, featuring the combined form of error function and its minimization process, offers the advantage of easy applicability for other researchers. Researchers can custom-design each error term according to their research needs and integrate it into the error function with an appropriate coefficient. While current experiments do not yet measure the activity of multiple neuron-class-identified neurons and the kymogram simultaneously, advancements in technology and the availability of such data will enable the creation of subjecting error terms. These terms will assess if the activities of neurons, beyond SMD neurons, accurately reproduce experimental observations. By minimizing the error function with these additional terms, a more advanced model can be developed in the future.

Our trained models also would enable the conduct of statistical experiments on characteristics not previously utilized in the creation of the error function. We offered 57 machine-learning trajectories that have learned both forward and backward locomotion, facilitating the statistical analysis of phenomena arising from researchers’ manipulation. For instance, to predict the effects of ablating neurons, one could ablate a specific neuron in each of the 57 machine-trajectories and statistically analyze the difference before and after the ablation to predict the effects of such ablation. This approach offers a mean for the preliminary testing of hypotheses that would otherwise be challenging or resource-intensive to explore through experimental methods.

### Limitations

Despite our efforts, our research still presents several limitations. First, the weight correlation in this study may have been overestimated. The distribution of anatomical and optimized synaptic weights does not converge into a single elliptical cluster (Fig. 2C, 2D). This implies the existence of a group that produced high correlation and another group that resulted in low correlation. Second, experimental findings(Yeon et al., 2018) indicated that *trp-1,2* mutation did not eliminate the oscillation of SMDD neuron activity, but merely altered its phase. However, *trp-1,2* mutation completely silenced the SMDD oscillation in the CANN model. Third, because the forward-locomotion-command and backward-locomotion-command were artificially set to constant values of 0 or 1, respectively, the activity of command neurons does not vary continuously, potentially diverging from the actual neuronal activity in *C. elegans*. Fourth, it is established that during forward (or backward) locomotion in *C. elegans*, the A-type (or B-type) motor neurons are not activate(Haspel et al., 2010). Conversely, in the CANN model, the membrane potential of A-type (or B-type) motor neurons oscillates under forward-locomotion-command (or backward-locomotion-command), presenting a notable discrepancy. To the best of our knowledge, no studies have yet addressed this discrepancy. Fifth, we utilized the anatomical connectome data from Cook et al., 2019, without questioning its reliability. However, this does not imply that we consider the data to be without any need for refinement. While we optimized the synaptic weights using our approach, we preserved the network topology, which could have led to the overestimation of anatomical tiny synapses during the optimization process.

## Acknowledgements

This work was supported in part by both pCoE program of DGIST [grant number: 23-CoE-BT-01] and a grant from iPT, Korea. TC and SK conceived the idea for this work. TC and SK set up the model, conducted the calculation, and analyzed the data and results. TC and SK wrote the manuscript. All authors reviewed and agreed on the contents of the manuscript. All data needed to evaluate the conclusions in the paper are present in the paper and the Supplementary Materials. The authors declare that they have no competing interests. We appreciate Prof. Iksoo Chang and Prof. Kyuhyung Kim for insightful discussions.

